# Correlation-based and feature-driven mutation signature analyses to identify genetic features associated with DNA mutagenic processes in cancer genomes

**DOI:** 10.1101/2020.01.06.895698

**Authors:** Hye Young Jeong, Jinseon Yoo, Hyunwoo Kim, Tae-Min Kim

**Affiliations:** Department of Medical Informatics, College of Medicine, The Catholic University of Korea, Seoul, Korea; Cancer Research Institute, College of Medicine, The Catholic University of Korea, Seoul, Korea; Department of Biomedicine and Health Sciences, Graduate School, The Catholic University of Korea, Seoul, Korea

## Abstract

Mutation signatures represent unique sequence footprints of somatic mutations resulting from specific DNA mutagenic and repair processes; however, their causal associations and potential utility for genome research remain largely unknown. In this study, we performed PanCancer-scale correlative analyses to identify the genomic features associated with tumor mutation burdens (TMB) and individual mutation signatures. We observed that TMB was correlated with tumor purity, ploidy, and the level of aneuploidy, as well as with the expression of cell proliferation-related genes representing genomic covariates in evaluating TMB. Correlative analyses of mutation signature levels with genes belonging to DNA damage-repair processes revealed that deficiencies of *NHEJ1* and *ALKBH3* may elevate TMB levels in cancer genomes accompanying APOBEC overactivity and DNA mismatch repair deficiency, respectively. We further employed a strategy to identify feature-driven, *de novo* mutation signatures and demonstrated they can be reconstructed using known causal features such as APOBEC overexpression, MLH1 underexpression, *POLE* mutations, and the level of homologous recombination deficiency. We further demonstrated, that tumor hypoxia-related mutation signatures are similar to those associated with APOBEC suggesting that APOBEC-related mutagenic activity mediates hypoxia-related mutational consequences in cancer genomes, and also, that mutation signatures can be further used to predict hypoxic tumors. Taken together, our study advances mutation signature-level mechanistic insights in cancer genomes, extending categories of cancer-relevant mutation signatures and their potential biological implications.

**Author summary:** Mutation signature analysis is powerful in deciphering the causative mutagenic events and their contributions in individual cancer genomes, but the causal relationship of individual mutation signatures are still largely unknown. PanCancer-scaled correlative analysis revealed mutation resource candidates in cancer genomes such as *NHEJ1* and *ALKBH3* deficiencies that may facilitate the accumulation of mutations in the setting of APOBEC overactivity and DNA mismatch repair deficiency, respectively. A feature-driven mutation discovery approach was employed to identify the mutation signatures representing homologous recombination deficiency and tumor hypoxia, the extent of which may serve as mutation-based phenotypic measures, previously estimated by DNA copy number alterations and mRNA expression signatures, respectively. Our study advances our understanding into the mechanistic insights of mutation signatures and proposes a method to utilize somatic mutations as a molecular proxy in terms of mutation signatures.

## Introduction

Recent advances in genomic sequencing technologies have yielded a huge catalog of somatic mutations in cancer genomes across diverse tumor types [1]. In addition to identifying cancer-driver mutations, including druggable targets [2, 3], clinical benefits associated with the quantitative nature of somatic mutations, such as the tumor mutation burden (TMB), have been demonstrated as predictive markers for immune checkpoint inhibitors [4–6]. The TMB of cancer genomes is highly variable within and between tumor types [7], with varying causal mechanisms that lead to hypermutation and mutator phenotypes [8]. To advance our understanding of the heterogeneity of TMB and its clinical relevance, it is essential to assess the roles of various exogenous and endogenous mutagenic agents and DNA repair-replication processes [9], as well as their relationship with genomic features.

Mutational processes are known to leave characteristic sequence features in genomes [10]. Well-recognized examples include C:G>A:T transversions and C:G>T:A transitions associated with tobacco smoke carcinogens in lung cancers [11] and ultraviolet (UV) radiation in skin cancers [12], respectively. The distinct sequence features of individual mutagenic processes indicate that mutational progressions that have been operative in cancer genomes can be inferred by sequence-based analyses. Accordingly, recently proposed trinucleotide context-based analysis of cancer genomes based on deconvolution techniques, such as non-negative matrix factorization (NMF), has revealed more than 30 mutation signatures across 7000 cancer genomes (hereafter, Sig1 to Sig30 as annotated in the Sanger mutation signature database; https://cancer.sanger.ac.uk/cosmic/signatures) [10, 13].

Signature-level mutation analysis enables the molecular dissection of TMB according to the distinct origins of the mutations because mutation signatures are often associated with causal genetic mechanisms or genes corresponding to endogenous mutagens and DNA repair-replication processes. For example, the overactivity of APOBEC cytidine deaminase leads to the accumulation of mutations consistent with Sig2 and Sig13, with sequence preferences of C>G and C>T within the Tp<u>Cp</u>N trinucleotide contexts [14]. Genetic events responsible for the deficiency of DNA repair enzymes have also been shown for some mutation signatures, e.g., promoter hypermethylation and transcriptional downregulation of *MLH1* [7] are responsible for the somatic DNA mismatch repair deficiency (MMRd) in colorectal and endometrial cancers leading to the accumulation of Sig6-consistent mutations. In addition, deregulation of DNA repair, or of the proofreading polymerase genes of *BRCA*-[15] and *POLE*-deficient genomes [16] leads to mutations consistent with Sig3 and Sig10, respectively [10]. However, the genetic mechanisms and potential gene markers which underlie the majority of mutation signatures are still largely unknown.

It is also possible that the current list of mutation signatures is not yet complete. Such missing mutation signatures may be found in somatic mutations with a unique presentation, *e.g.,* a recent mutation signature analysis focused on clustered mutations revealed a novel signature associated with the activity of a translesional polymerase of *POLH* [17]. We also reported a cisplatin treatment-related mutation signature that was exclusively observed in head and neck cancers with a history of chemotherapy, not in treatment-naïve cancer genomes [18]. Given that the current list of mutation signatures has been largely obtained by unsupervised deconvolution-based methods, such as NMF, methods using differential nucleotide frequencies may lead to the identification of novel mutation signatures representing specific tumor phenotypes or genotypes.

Large-scale multiomics cancer genome data may serve as valuable resources for identifying underlying genetic mechanisms and key DNA repair genes that contribute to somatic mutations and the TMB levels of cancer genomes. In this study, we performed PanCancer-scaled correlative analyses that linked TMB and mutation signatures with multiomics datasets available from the PanCancer-scale Cancer Genome Atlas (TCGA) consortium. We first evaluated the correlation of TMBs with systematic genomic features, such as tumor purity, ploidy and aneuploidy. Then, TMB was deconvoluted into the levels of known mutation signatures (i.e., the degree to which Sig1 to Sig30 individually contributed in given cancer genomes), which were subject to multiomics correlative analyses to discover known and novel relationships between mutation signatures and their potential genetic mechanisms. We also propose a feature-driven mutation signature discovery method to identify *de novo* mutation signatures as differential trinucleotide frequencies based on tumor features of interests, such as homologous recombination (HR) deficiency or tumor hypoxia.

## Results

### Genomic features associated with TMB across cancer genomes

To identify potential genomic covariates of TMB, we performed a PanCancer-scale correlative analysis with systematic genomic features, such as tumor purity and ploidy for 9,857 TCGA cases. First, we observed that the log-transformed TMB was inversely correlated with tumor purity (*r* = -0.118; **Figs 1A** and **1B**). We previously showed that tumor purity represents a surrogate marker for the level of tumor-infiltrating immune cells, including cytotoxic T cells [19], and the observed purity-TMB relationship may reflect an association between the level of tumor-infiltrating immune cells and TMB. We also observed that TMB was positively correlated with the level of aneuploidy, i.e., TMB was correlated with tumor ploidy (*r* = 0.161; **Figs 1C** and **1D**) and also with the number of dosage-imbalanced copy number segments (*r* = 0.291; **Figs. 1E** and **1F**). We also performed gene set enrichment analysis (GSEA) to identify molecular functions associated with the TMB (**S1** and **S2 Tables**). Genes whose expression levels are positively or inversely correlated with TMB were enriched in cell-cycle and ion transport functions, respectively. Cell cycle-related replication stress is known to be a potential cause of mutations [20] and the transcriptional activation of cell-cycle-related genes may be related to the number of cell cycles in cancer stem cells. However, given that the TMB represents an admixture of mutations resulting from varying mutagenic or repair processes, the correlative analysis of TMB is limited in identifying the specific causal genes or mechanisms.

**Fig 1.**
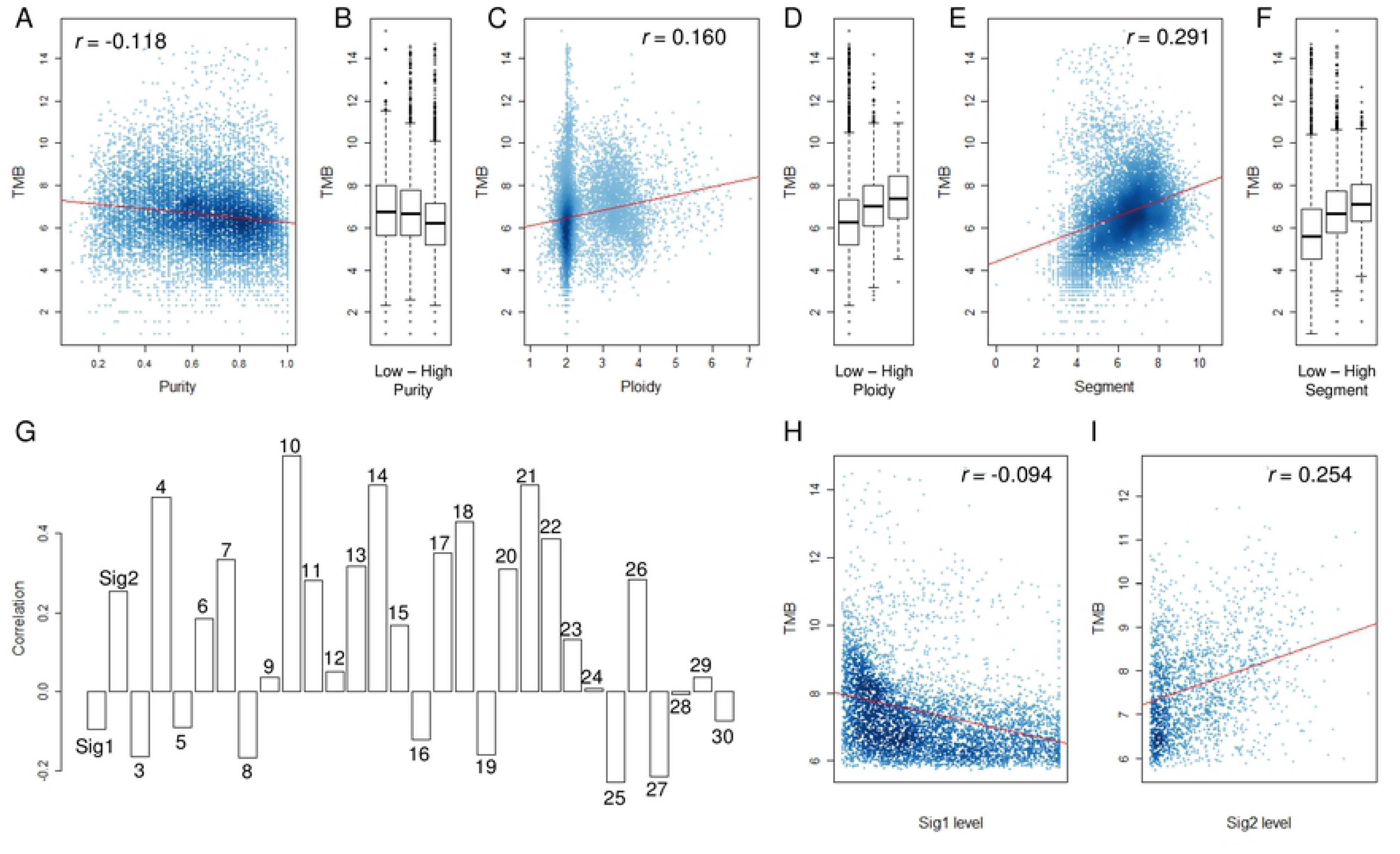
The relationship between TMB and other genomic features. (A) A scatter plot shows the inverse correlation between log2-transformed tumor mutation burden (TMB) and tumor purity. (B) TMB is shown for three equal-sized tumor bins, *i.e.,* low-moderate-high tumor purity. (C and D) The positive correlation between TMB and tumor ploidy levels. (E and F) The positive correlation between TMB and the number of dosage-imbalanced segments. (G) Correlation level between TMB and the 30 mutation signature levels (S1-S30). (H and I) The correlation between TMB and level of Sig1 and Sig2 are shown in scatter plots as examples of inverse and negative correlation with TMB.

The deconvolution-based, mutation signature levels (*i.e.,* the relative abundance of mutations that belong to the corresponding mutation signatures) represent the relative contribution and activity levels of specific mutagenic or repair processes and represent a more appropriate resource for the correlative analyses. Accordingly, we obtained 30 mutation signatures (Sanger ver. 2 mutation signatures, annotated Sig1-Sig30). For PanCancer tumor specimens, we estimated the relative contribution or signature levels of all 30 [21] and performed correlative analyses with TMB (**Fig 1G**). Notably, the correlations between individual mutation signatures and TMB were variable, suggesting that not all signatures are responsible for the emergence of hypermutation in cancer genomes. For example, inverse and positive correlations with TMB were observed for Sig1 and Sig2 levels, respectively (**Figs 1H** and **1I)**. The Sig1-associated mutations that have accumulated continuously during the lifetime of a host before the malignant transformation, are minimally associated with hypermutation or the acquisition of mutator phenotype in cancer genomes. The enzymatic activity of APOBEC cytidine deaminase, which is responsible for the generation of Sig2 mutations, can be variable often leading to hypermutations. A recent study demonstrated that APOBEC activity may be associated with episodic mutation bursts in cancer cell lines [22]. Correlation levels with TMB were relatively high for mutation signatures associated with well-recognized causal events (e.g., Sig2/APOBEC, Sig4/smoking, Sig6/MMRd, Sig7/UV, and Sig10/POLE), suggesting these events can contribute to the high mutation load of cancer genomes. Scatterplots showing the correlation levels for 30 mutation signatures are shown in **S1 Fig**.

### Mutation signature correlative analyses for DNA damage and repair genes

To assess the feasibility of exploiting multiomics datasets containing expression data, to support previously established relationships, we first examined *MLH1,* whose promoter methylation, along with transcriptional downregulation leads to MMRd in microsatellite instability-high (MSI-H) genomes [23]. Positive and inverse correlations were consistently observed between *MLH1* expression and TMB (*r* = 0.339) and between *MLH1* promoter methylation levels and TMB (*r* = -0.349) where the relationship is largely attributed to the presence of MSI-H genomes (red dots; **S2 Fig**).

We next estimated the relative contribution or signature levels of 30 known mutation signatures [21] and performed correlative analyses with expression levels of 254 genes belonging to the DNA damage and repair (DDR) pathway [24]. The distribution of PanCancer-scale correlation levels are shown for gene expression and promoter methylation levels of the DDR genes in **Figs 2A** and **2B**, respectively. As expected, the lowest level of correlation with gene expression of *MLH1* (*r* = -0.551; an arrow in **Fig 2A**) and the second-highest level of correlation with promoter methylation of *MLH1* (*r* = 0.233; an arrow in **Fig 2B**) were observed with Sig6 levels representing MMRd. The correlation levels for individual mutation signatures with DDR gene expression and promoter methylation are available in **S3 S4 Tables**, respectively.

**Fig 2.**
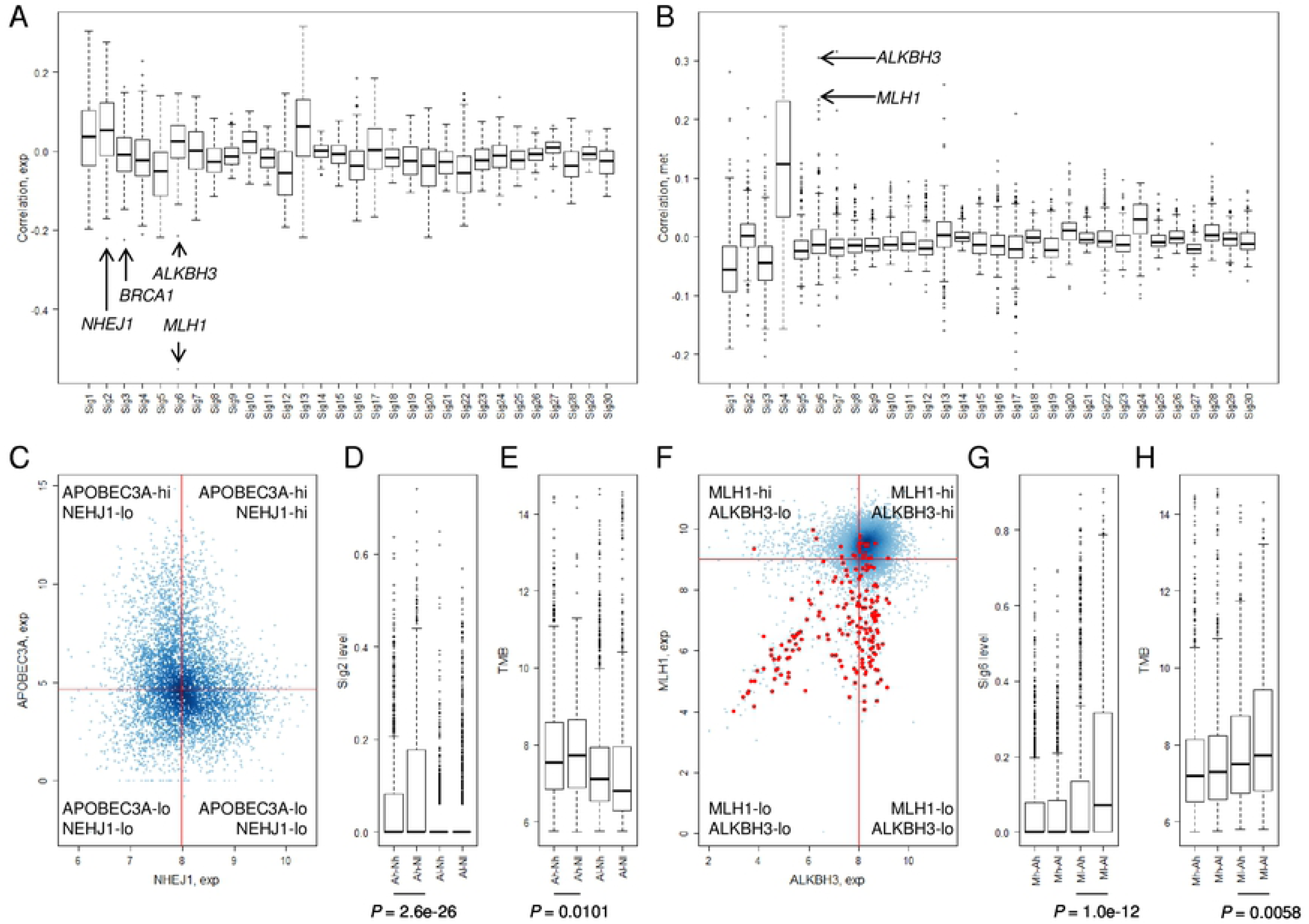
Correlative analysis of mutation levels with the DDR gene. (A) The levels of 30 mutation signatures were correlated with expression of 254 DDR genes. The *Y*-axis shows correlation level, with arrows for selected genes (*ALKBH3*, *BRCA1*, *MLH1*, and *NHEJ1*). (B) The correlation between the 30 mutation signature levels and DDR gene promoter methylation levels are shown. (C) Scatter plot shows the expression level of *APOBEC3A* and *NHEJ1* (log-scaled). The cases were discriminated into four classes using the medians of *APOBEC3A* and *NHEJ1* expression (shown by red lines). (D) A significant difference was observed in the Sig2 levels of tumors with *APOBEC3A* overexpression according to *NHEJ1* expression ([A]POBEC3A[h]igh-[N]HEJ1[h]igh-*vs.*-Ah-Nl, *P* = 2.6e-26; *t*-test). (E) TMB levels were also significantly different between Ah-Nh-*vs.*-Ah-Nl (*P* = 0.01; *t*-test). (F) Scatter plot show the distribution of *ALKBH3* expression (*x*-axis; log2-scaled) and *MLH1* expression (*y*-axis; log2-scaled). Red dots indicate MSI-H cases. (G and H) A significant difference was observed for Sig6 levels and TMB levels for tumors with down-regulated *MLH1* according to *ALKBH3* expression levels ([M]LH1[l]ow-[A]LKBH3[h]igh-*vs.*-MlAl; *P* = 1.0e-12 and *P* = 0.0058 for Sig6 levels and TMB levels, respectively; *t*-test).

In addition to *MLH1*, a substantial level of inverse correlations were observed for certain DDR gene expression and mutation signature pairs, e.g. *NHEJ1*-Sig2 (*r* = -0.221; 1^st^ ranked in Sig2, **Fig 2A**) and *BRCA1*-Sig3 (*r* = -0.224; 1^st^ ranked in Sig.#3, **Fig 2A**), suggesting that their deficiency may give rise to mutations belonging to the corresponding mutation signatures. The association between *BRCA* loss and Sig3 representing HR deficiency has been well documented [15]. However, the association between *NHEJ1,* that encodes essential DNA repair factors mediating non-homologous end-joining, with Sig2 is not well known. Furthermore, *NHEJ1* expression was also negatively associated with the level of Sig13 (*r* = - 0.212; 1^st^ ranked in Sig13; **Fig 2A**), which is similar to Sig2 in potential causality and nucleotide composition [10]. Hierarchical clustering of joint profiles of mutation signature levels and DDR gene expression also highlights the association between *NHEJ1* and Sig2-Sig13 (**S3 Fig**). Since no substantial correlation between *NHEJ1* and *APOBEC3A* was noted, we classified the genomes into four classes according to the median expression of these two genes (*<u>A</u>POBEC3A*-<u>hi</u>gh/<u>lo</u>w and *<u>N</u>HEJ1*-<u>hi</u>gh/<u>lo</u>w; **Fig 2C**) and the Sig2 levels are shown against the four classes (**Fig 2D**). Genomes with low expression levels of *APOBEC3A* were almost devoid of Sig2 levels, regardless of *NHEJ1* expression levels. However, the *APOBEC3A* up-regulated genomes were further discriminated into two classes according to *NHEJ1* expression levels with significant differences in the Sig2 levels (A-hi, N-hi *vs.* A-hi, N-lo*, P* = 2.6e-26; *t*-test). These findings suggest that *NEHJ1* deficiency alone does not contribute to Sig2 mutations without *APOBEC3A* activity; however, an *NHEJ1* deficiency may potentiate the mutagenic activity of APOBEC cytidine deaminase. Given the roles of *NHEJ1* in double-strand breakage (DSB) repair [25], it is assumed that an *NHEJ1* deficiency may delay the repair of DSB leaving the transient single-strand terminus open as a substrate for APOBEC mutagenesis also leading to an elevated TMB that is significant (*P* = 0.0101, *t*-test; **Fig 2E**).

For Sig6 associated with *MLH1* deficiency and MMRd, we also observed substantial correlation with *ALKBH3* expression and *ALKHB3* promoter methylation (*r* = -0.215 and *r* = 0.304; 2^nd^ and 1^st^ ranked in Sig6, respectively; **Figs 2A** and **2B**). The relationship between the methylation levels of *MLH1* and *ALKBH3* and Sig6 levels are illustrated in the hierarchical clustering of the joint methylation profiles and mutation signature levels (**S4 Fig**), as well as those of the DDR gene expression profiles (**S3 Fig**). The prevalent epigenetic modification of *ALKBH3* has been recently reported [24] but its functional significance is largely unknown. A similar regulatory relationship such as the inverse correlation between the expression and methylation of *MLH1* and *ALKBH3* is not simply explained by genomic adjacency (*ALKBH3* on 11p11.2 and *MLH1* on 3p22.2, respectively). The similar regulatory relationship of *ALKBH3* and *MLH1,* as well as their substantial correlation with the MMRd representing mutation signature, suggests that *ALKBH3* deficiency may help the accumulation of Sig6-consistent mutations in the context of MMRd induced by *MLH1* deficiency. We observed that MSI-H genomes consistently show down-regulation of *MLH1* (red dots in **Fig 2F**) but MSI-H genomes can be further classified into those with or without down-regulation of *ALKBH3* (**Fig 2F**). In addition, significant differences in Sig6 levels and TMB were observed between genomes with or without *ALKBH3* down-regulation (*P* = 1.0e-12 and *P* = 0.0058, respectively).

### Feature-driven discovery of mutation signatures

The correlation of gene expression or other genomic features such as DNA promoter methylation levels, with their attributed mutation signature levels, suggests that the corresponding mutation signatures can be directly inferred using genotypic or phenotypic class distinction. For example, *de novo* mutation signatures representing a deficiency in DDR genes can be derived as differential trinucleotide frequencies between genomes with high and low expression of the gene. We tested the method for three genes whose transcript levels or somatic mutations were associated with the cognate mutation signatures – Sig2/*APOBEC3A* (high-expression), Sig6/*MLH1* (low-expression), and Sig10/*POLE* (somatic mutation) (**Fig 3**). The mutation signature representing APOBEC overactivity was derived as differential trinucleotide frequencies between genomes with high expression of *APOBEC3A* and those with low expression as shown in **Fig 3A**. The MMRd-representing signature was also inferred from the comparison of genomes with low and high *MLH1* expression. Genomes with *POLE* mutations were also compared to those without *POLE* mutations to derive *POLE*-related mutation signatures. Since the negative contribution of mutation profiles is not biologically meaningful in terms of mutation signatures, only positive differentials were taken into account. Thus, only positive differentials were taken for *APOBEC3A* (high-expression) minus *APOBEC3A* (low-expression) with the direction indicated (e.g. *APOBEC3A*-expression high). **Fig 3B** shows three mutation signatures derived from 96 trinucleotide frequencies, along with their cognate mutation signatures (Sig2, Sig6 and Sig10 matching with *APOBEC3A*/high-exp, *MLH1*/low-exp and *POLE*/mutated, respectively). **Fig 3C** also shows that feature-driven mutation signatures were segregated along with their cognate signatures in terms of trinucleotide frequencies suggesting that the inferred mutation signatures based on the genotypic and phenotypic class distinction can mimic their cognate mutation signatures.

**Fig 3.**
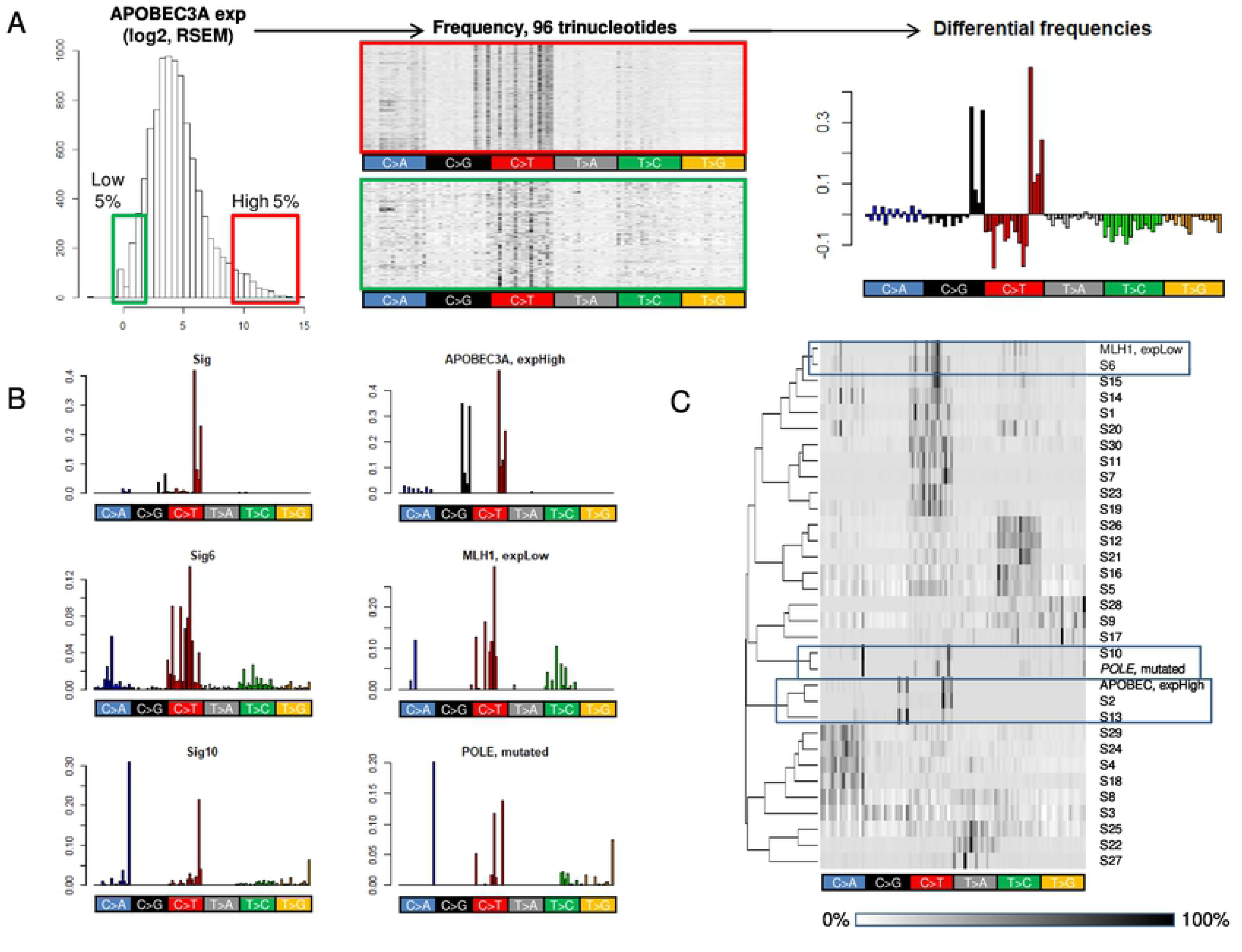
Feature-driven mutation signatures. (A) Histogram show the distribution of expression of *APOBEC3A* (log2 scale) with tumors up-regulated for *APOBEC3A* (5 percentiles, red box) and down-regulated (5 percentiles, green box) selected (left). Heatmaps showing the 96 cases with trinucleotide frequencies of APOBEC3A up- and down-regulated, (red and green boxes, respectively; middle). The differential of average nucleotide frequencies (APOBEC3A up-*vs*. down-regulated) is shown as an *APOBEC3A* expression-driven mutation signature (right). (B) Bar plots showing the trinucleotide frequencies of Sig2, Sig6, and Sig10 (left). The differentials of the trinucleotide frequencies are shown as feature-driven mutation signatures. *APOBEC3A* expression (high-*vs.*-low), *MLH1* expression (low-*vs.*-high), and *POLE* mutations (*POLE* mutant-*vs.*-wild types) were used as features to derive mutation signatures showing resemblance to the cognate mutation signatures (Sig2, Sig6 and Sig10, respectively). In the *APOBEC3A expression* example, we selected the positive differential values as ‘APOBEC3A-highExp’ for subsequent analysis (right). (C) Hierarchical clustering shows that Sig2, Sig6, and Sig10 are segregated with mutation signatures derived from *APOBEC3A* expression, *MLH1* expression, and *POLE* mutations, respectively.

### Mutation signatures representing HR deficiency

We further tested whether genomic features other than single gene activities could be used for the discovery of *de novo* mutation signatures. Thus, we obtained three somatic copy number alteration (SCNA)-based scores representing HR deficiency (NtAI, LST, and HRD-LOH) for TCGA tumors [26]. As expected, the three HR deficiency score-driven mutation signatures were similar to Sig3 in terms of trinucleotide frequencies (**Fig 4A**). Hierarchical clustering of 96 trinucleotide frequencies also segregated those of 3 HR deficiency-driven mutation signatures along with Sig3, further highlighting the trinucleotide context-level similarities (**Fig 4B**). Next, we explored whether the mutation signature levels were correlated with the original SCNA-based scores of HR deficiency (**Fig 4C**). The mutation signature levels were estimated by metagene projection for each HR deficiency score and then were correlated with the original score. Substantial correlation levels (*r* = 0.465 – 0.499) were observed, suggesting that the SCNA-based HR deficiency scores could be reproduced to some extent, from somatic mutations in terms of the mutation signatures (**S4 Fig** for individual correlations). SCNAs and mutations are distinct types of genomic alterations in cancer genomes and it is still controversial whether the abundance of SNCAs and mutations are simply correlating with each other [27, 28]. Our findings suggest that genomic events (e.g., HR deficiency) may leave genomic footprints across different aspects of cancer genomes such as SCNAs and somatic mutations. Once we identify the mutation signatures corresponding to the genomic events, the level of contribution in given genomes can be estimated by signature refitting or metagene projection as we propose. The good concordance level between SCNA-based HR deficiency scores and the inferred mutation signature levels highlight the potential utility of somatic mutations as genomic markers

**Fig 4.**
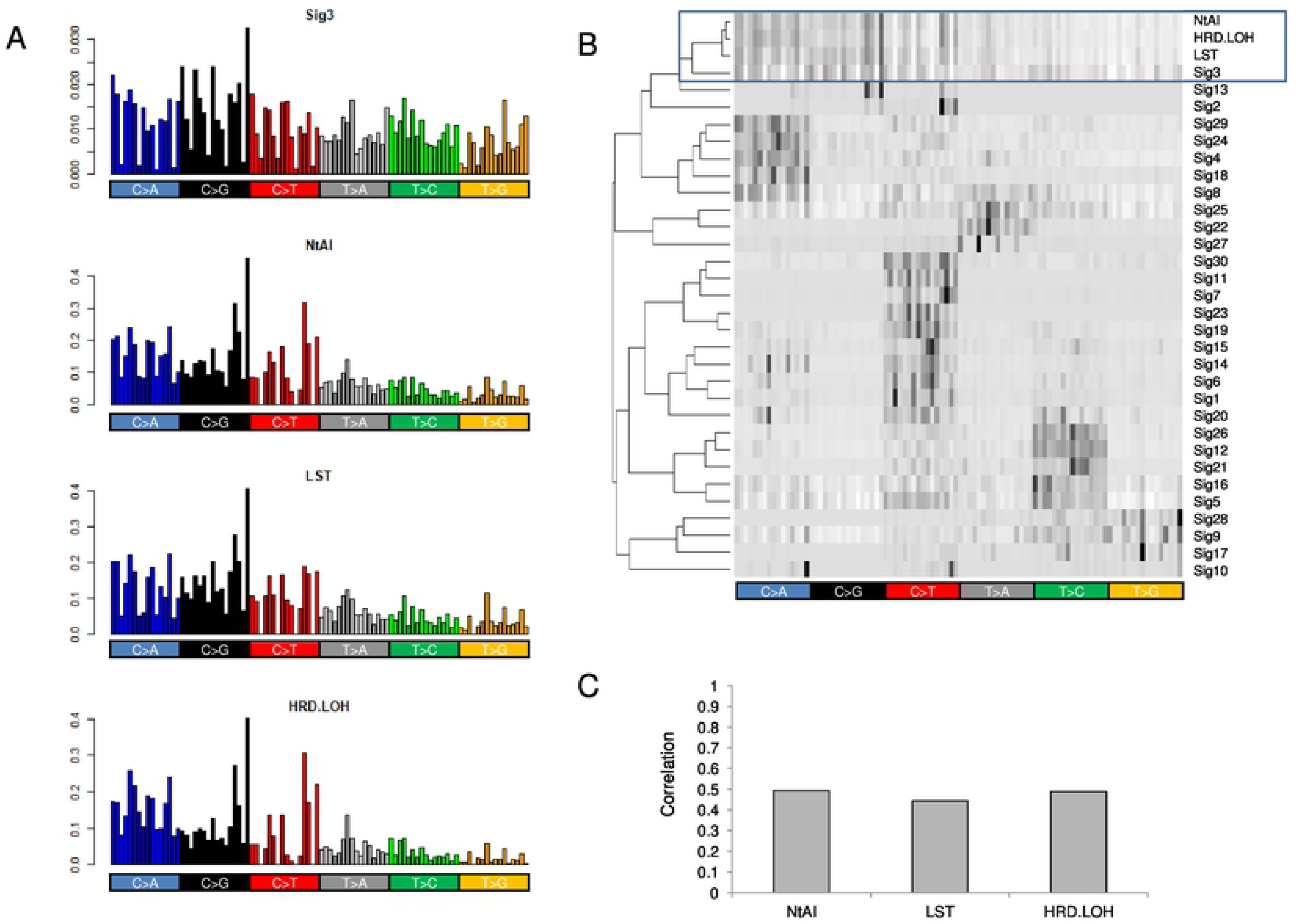
Mutation signatures of HR deficiency. (A) Sig3 shows similar trinucleotide frequencies distribution with those inferred from the three HR deficiency scores of NtAI, LST and HRD-LOH. (B) Heatmap showing that three mutation signatures inferred by HR deficiency scores are segregated with the Sig3 mutation signature. (C) Correlation levels are shown for SCNA-based HR deficiency scores and mutation signature levels estimated by metagene projection.

### Mutation signatures representing tumor hypoxia

To further test feature-driven mutation signatures, we selected tumor hypoxia as one of the key tumor hallmarks associated with poor prognosis and treatment failure [29]. We obtained eight mRNA signature-based tumor hypoxia scores of TCGA tumors from the literature [29]. The mutation signatures of tumor hypoxia were then identified as differential trinucleotide frequencies of mutations between hypoxic and normoxic tumors (*i.e.,* tumors with high and low mRNA-based hypoxia scores) for each of eight hypoxia scores. Of note, hierarchical clustering showed that six of the eight hypoxia score-based mutation signatures were co-segregated with APOBEC-related Sig2 and Sig13 (red open box, **Fig 5A**)., The hypoxic tumors identified by the mRNA-based hypoxia scores consistently showed elevated Sig2 levels compared to normoxic tumors for six out of 8 hypoxia scores, four of which were significant (**Fig 5B**). This suggests that the genomic consequences of tumor hypoxia, at least for somatic mutations, can be substantially attributed to APOBEC enzymatic activity. We further performed a test to predict hypoxic tumors based solely on the mutation signatures. Mutation signature levels of individual tumors were estimated by metagene projection and the resulting scores were used to call the mutation signature-based hypoxic tumors. Calls made by mRNA-based hypoxic scores were compared to examine the concordance with mutation signature. We observed that concordance was >75% for seven hypoxic signatures (**Fig 5C**). These findings suggest that the impact of hypoxia in terms of somatic mutations may have specific nucleotide predisposition and that tumor hypoxia may be predicted by the mutation profiles of cancer genomes. Again, our results show that the somatic mutations in terms of mutation signatures can be used as genomic markers and the inferred signature levels as quantitative measures showed good concordance to tranditional markers such as SCNA-based HR deficiency scores and mRNA-based tumor hypoxia scores.

**Fig 5.**
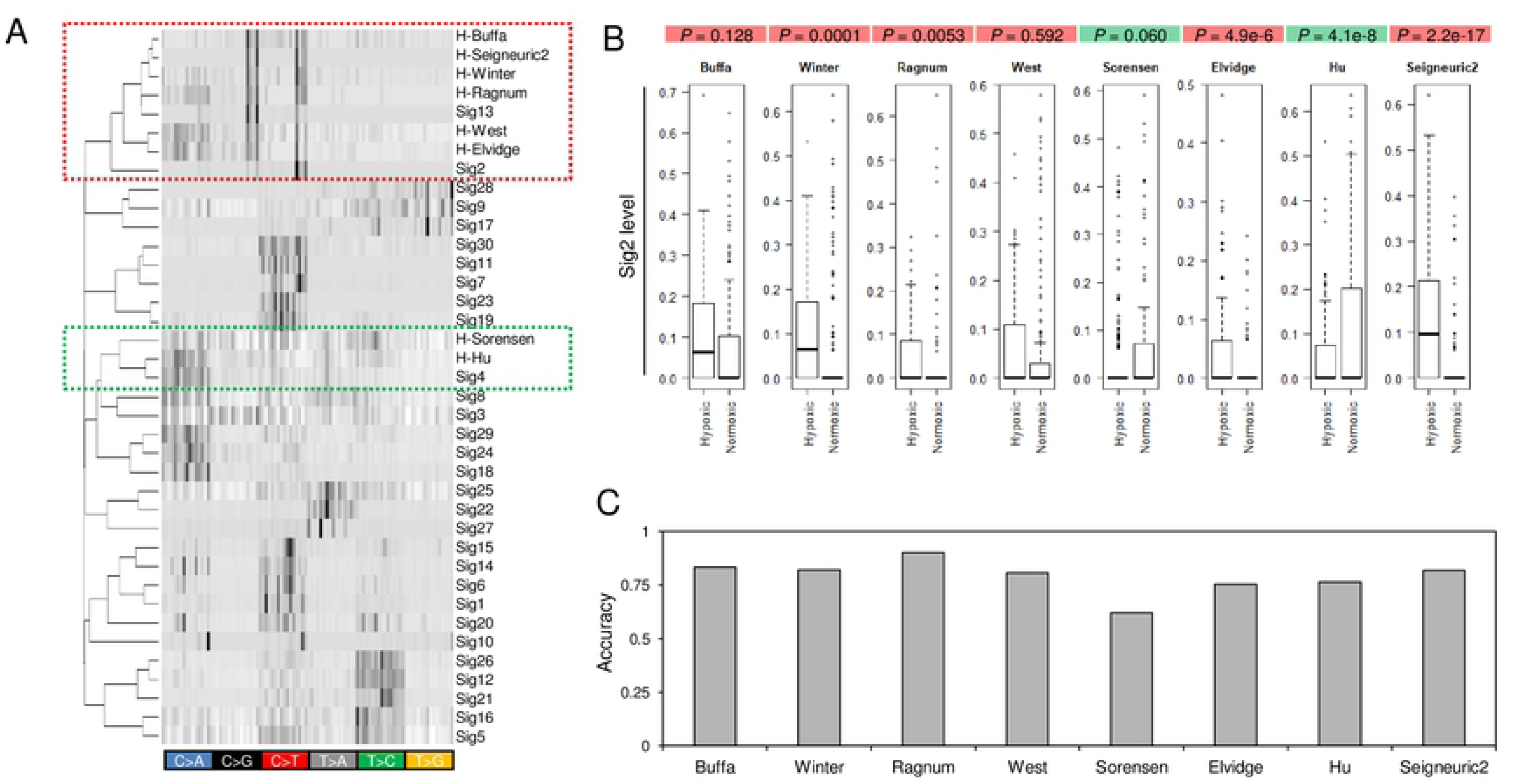
Mutation signatures of tumor hypoxia. (A) Eight mutation signatures were derived based on the mRNA-based tumor hypoxia scores with literature-based annotation as obtained. A heatmap is shown to demonstrate that six out of eight tumor hypoxia-representing mutation signatures are co-segregated with APOBEC-related Sig2 and Sig13 (red open box, upper) for 96 trinucleotide contexts. Two hypoxia mutations were also co-segregated with Sig4 (green open box, below). (B) Boxplots showing Sig2 levels for hypoxic and normoxic tumors as annotated according to the mRNA-based hypoxic scores. The significance of difference of Sig2 levels between hypoxic and normoxic tumors are shown at top (*t*-test). Six hypoxic scores segregated with Sig2/Sig13 were shown with red while those with normoxic tumors are green. (C) Accuracy is shown across eight hypoxia scores as the level of concordance between the hypoxia-normoxia calls made by inferred mutation signature levels and mRNA score-based calls.

## Discussion

In this study, we performed PanCancer-scaled correlative analyses for the TMB and their deconvoluted mutation signatures with various genomic features, including the expression and methylation of DDR genes. We further proposed an analytical framework to derive feature-driven mutation signatures based on genotypic and phenotypic class distinctions. TMB, a measure of the total number of mutations in a given cancer genome, has been recently highlighted as a useful biomarker for treatment with immune checkpoint inhibitors [5, 30]. This study demonstrated that systematic genomic variables of cancer genomes, such as tumor purity, ploidy, and the level of aneuploidy, are correlated with TMB. The observed correlation may indicate a causal relationship between variables but it is also possible that the identified genomic features represent confounding factors. For the latter, the correlating features can be taken into account in TMB-centric correlative analysis. An example of this being the assessment of clinical benefits for immunotherapy in cases with high TMB, as we have recently demonstrated that tumor purity was a confounding factor for cancer genome analyses [19]. Some mechanistic insights can be inferred from the correlation results, *e.g.,* the overall positive correlation between TMB and the level of aneuploidy may indicate that chromosome-(ploidy and aneuploidy levels) and mutation-level instability (TMB) are related to each other in cancer genomes. But, the pathway analysis of GSEA in search of specific molecular functions associated with TMB only revealed that the universal cellular functions of cell cycling and chromosome-related genes were relatively overexpressed in high-TMB tumors. This is consistent with a previous assumption that the number of cell cycles and thus, elevated cell cycling in cancer stem cells, may be associated with elevated TMB [20]. However, since TMB of individual cancer genomes represents the aggregate of multiple mutagenic and DNA repair-replicative processes, TMB-based correlative analysis hardly points to specific DNA mutagenic or repair/replicative processes with potential biological or clinical relevance. To cope with this issue, we deconvoluted the TMB into known, multiple mutation signatures and used their levels for the correlative analyses. We focused on the expression and DNA promoter methylation of 254 DDR genes belonging to nine DNA damage-repair-replicative processes [24]. Along with *MLH1,* whose promoter hypermethylation and resulting transcriptional downregulation leads to somatic MSI-H genotypes and the generation of mutations belonging to Sig6, we observed an additional relationship between Sig2-*vs.*-*NHEJ1* deficiency and Sig6-*vs.*-*ALKBH3* deficiency. In both cases, deficiencies in *NHEJ1* and *ALKBH3* alone did not increase mutations corresponding to cognate signatures or TMB. Instead, their deficiency is effective only in genomes with APOBEC overactivity and MMRd, respectively. This raises an assumption that *NHEJ1* deficiency supplements APOBEC-related mutagenesis, or the deficiency of *ALKBH3* with roles in alkylation damage reversal may synergize with the deficiency of other DNA repair classes such as MMRd. These hypotheses need further validation and the list of DDR genes showing high levels of correlation may serve as potential candidates in the search of hypermutated cancer genomes, with clinical benefits [8].

Currently, available mutation signature discovery methods are classified into two categories, i.e., those for “*de novo*” mutation signature discovery and the others for “signature refitting” using known mutation signatures [31]. For the former, unsupervised NMF or its derivatives have been proposed to extract the *de novo* mutation signatures whose lineage specificity and potential causal association are investigated *post hoc*. However, it has rarely been discussed whether mutation signatures can be derived in a supervised manner directly from phenotypic or genotypic scores of interest, and can serve as a proxy or marker for genomic studies. To test the viability of feature-driven discovery of mutation signatures, we first demonstrated that Sig2, Sig6, and Sig10, which are known to be associated with *APOBEC* over-expression, *MLH1* under-expression, and *POLE* mutations, respectively, could be derived using the associated genetic features, that is the differential trinucleotide frequencies between genomes with or without the known causal events. Next, we used three SCNA-based scores representing HR deficiency to derive mutation signatures, which were similar to Sig3 associated with *BRCA* deficiency. The mutation signatures inferred from the HR deficiency scores showed similarities with Sig3 in terms of 96 trinucleotide contexts, and more importantly, the mutation signature levels estimated from mutation profiles showed a good concordance to the original SCNA-based HR deficiency scores. This suggests that the mutation profiles could be used to infer the level of HR deficiency, primarily done with SCNA profiles. Although the use of quantitative features of somatic mutations has been largely limited to TMB, our study further demonstrated that somatic mutations in terms of mutation signatures could serve as cancer markers. Finally, tumor hypoxia has been proposed to increase mutation rates of cancer genomes with the downregulation of MMR genes [32], and hypoxia is also associated with HR, *TP53* and *RAD53* deficiencies [29, 33, 34]; however, the impact of tumor hypoxia on tumor mutations is not well understood. This study showed that the majority of tumor hypoxia-driven mutation signatures resembled those of the APOBEC-related signatures of Sig2 and Sig13 in terms of trinucleotide frequencies. Along with the elevated Sig2 levels in hypoxic tumors, this suggests that tumor hypoxia is associated with APOBEC activity and that the somatic mutations in hypoxic tumor genomes may be largely attributed to the APOBEC-mediated C-to-T transitions. We further demonstrated that the mutation signature levels inferred from the mutation profiles of cancer genomes can be used to predict the hypoxic tumors. Again, this indicates that the mutation profiles can be used as proxies to infer various cancer genome-related features by mutation signature-based analysis. Although our exploratory study requires further validation in an extended, validation cohort, the potential of mutation signatures to derive cancer hallmark features, such as HR deficiency and tumor hypoxia, may be extended to mutation signature-based marker discovery with potential clinical relevance.

## Materials and methods

### Mutation information

The TCGA cancer mutation profiles as well as other multiomics datasets encompassing 9,587 tumor specimens and > 30 tumor types were downloaded from the TCGA PanCancer Atlas (https://gdc.cancer.gov/about-data/publications/pancanatlas). Duplicate cases were removed and only mutations from primary tumor genomes were considered, ignoring those from recurrent or metastatic genomes. TMB was defined as the sum of all types of somatic calls for single nucleotide variations and short indels. We also obtained the tumor purity and ploidy data, as well as other copy number-related variables from the literature [35]. For SCNA profiles, we obtained segment profiles of primary tumors from the TCGA PanCancer Atlas. The copy number imbalanced segments were defined as those with log2 ratio > 0.2 or <-0.2 and the sum of segment size was normalized by genome size to obtain genomic fraction (%). MSI-H cases were those annotated by Bethesda marker-based tests for four tumor types of colorectal, stomach, endometrial and esophageal cancers [36].

### GSEA

To identify molecular functions associated with TMB, we calculated Pearson’s correlation coefficients for individual gene expressions (log-scaled RSEM) with TMB (log-scaled). Gene-level correlations were subjected to the pre-ranked version of gene set enrichment analysis (GSEA) with Gene Ontology (GO) terms [37] available in MSigDB (http://software.broadinstitute.org/gsea/msigdb/index.jsp; c5 category).

### Signature analysis

For known mutation signatures, we download 30 signatures as Sanger ver.2 mutation signatures (Sig1 – Sig30, https://cancer.sanger.ac.uk/cosmic/signatures). For signature refitting, we used deconstructSigs R packages [21] to derive the relative contribution of individual mutation signatures in given cancer genomes. The estimated contribution or the number of mutations belonging to individual signatures were used as mutation signature levels for the subsequent correlative analyses. For mutation signature analysis, we used 6,040 cancer genomes harboring no less than 50 mutations. For correlation with mutation signature levels, we selected 254 genes that belong to DNA damage and repair (DDR) processes available in a previous study [24]. The transcriptome sequencing-based expression levels and microarray-based DNA promoter methylation levels of DDR gene were also obtained from the TCGA PanCancer Atlas.

### Feature-driven mutation signature analysis

A mutation signature representing APOBEC overactivity was derived as the differentials of trinucleotide frequencies between tumors with high and low expression of *APOBEC3A* (95th and 5th percentiles, respectively). For mutation signatures representing *MLH1* deficiency, the differentials were calculated between expression levels of *MLH1*. Only positive values of differential trinucleotide frequencies were considered, given that negative values have no biological significance for signature-based analysis. Thus, ‘APOBEC3A-high expression’ signatures are the positive values when looking at diffentials of APOBEC3A high – APOBEC3A low tumors and represent APOBEC overactivity. ‘MLH1-low expression’ signatures were similarly generated to represent *MLH1* deficiency. For the *POLE*-signature, the differential of trinucleotide frequencies were obtained from comparisons of *POLE*-mutated and wild-type genomes. Such feature-driven mutation signatures were compared with known mutation signatures (Sig1-Sig30) using hierarchical clustering of the trinucleotide frequencies.

### Mutation signatures representing HR deficiency and tumor hypoxia

Three HR deficiency-representing scores representing telomeric allelic imbalance (NtAI), large scale transition (LST), and loss of heterozygosity (HRD-LOH), were obtained from a previous publication [26]. Eight scores of tumor hypoxia estimated from mRNA signatures were also obtained from the literature [29]. Two types of mutation signatures were acquired per score. For an example of a hypoxia score, positive and negative differential values of the trinucleotide frequencies between genomes with high and low hypoxia scores were obtained as hypoxia-and normoxia-representative mutation signatures. To estimate the mutation signature levels using two signatures for each hypoxia score, we employed metagene projection, where the positive linear combination of the two mutation signatures, *i.e.,* hypoxia and normoxia signatures from a single score, were projected onto the normalized mutation frequencies across the genomes to be examined [38]. For metagene projection, we used Moore-Penrose generalized pseudoinverse with the ginv function of the R MASS library as previously described [39]. For HR deficiency-related mutation signatures, we calculated the correlation level between the mutation signature levels representing tumor hypoxia and the corresponding SCNA-based HR deficiency scores as the concordance levels. For tumor hypoxia, the signature levels representing tumor hypoxia and normoxia (by calculating correlation levels with Sig2 and binary calls of ‘hypoxia’ or ‘normoxia’) were made for individual tumors. The mutation signature-based calls were compared with mRNA score-based calls (e.g., hypoxic and normoxic tumors) and the concordance levels were used to assess accuracy.

## Acknowledgements

We appreciate the kind support during manuscript preparation of members of the Cancer Research Institute, Catholic University of Korea.

## Supporting information

**S1 Fig. Correlation between mutation signature levels and TMB.** Scatter plots show the correlation levels between 30 mutation signatures (Sig1 to Sig30) and TMB.

**S2 Fig. Relationship between MLH1 expression-methylation levels with TMB.** (A) TMBs are shown against the expression level of *MLH1*. Red dots indicate tumors with high microsatellite instability-high (MSI-H). (B) Similarly shown for the level of promoter methylation of *MLH1*.

**S3 Fig. Clustering of joint profiles of DDR gene expression and mutation signature levels.** (A) The expression level of 254 DDR genes were merged with those of mutation signature levels estimated from 5617 TCGA samples and were subjected to hierarchical clustering. Inset shows that *NHEJ1* expression was segregated with the level of Sig2 and Sig13 associated with APOBEC overactivity. (B) Scatter plots show *NHEJ1* expression levels and the levels of Sig2 and Sig13, respectively.

**S4 Fig. Clustering of joint profiles of DDR gene methylation and mutation signature levels.** (A) The promoter methylation levels of 254 DDR genes were merged with those of mutation signature levels and were subjected to hierarchical clustering. Inlet shows that the methylation levels of *MLH1* and *ALKBH3* segregated with the level of Sig6. (B) Scatter plots show the level of correlation of gene methylation (x-axis) and expression (y-axis) with the level of Sig6, highlighting the *MLH1* and *ALKBH3* deficiencies. (C) The correlation of *MLH1* expression and methylation levels are shown in a scatter plot (red-dots for MSI-H cases). (D) Similarly shown for *ALKBH3*.

**S5 Fig. Concordance of known and feature-driven mutation signature levels.** For three HR deficiency scores, mutation signatures were identified and the signature levels were estimated by metagene projection (x-axis). Y-axis shows the known signature levels of Sig3 representing *BRCA* or HR deficiency.

**S1 Table. Molecular functions enriched for high TMB.** Preranked GSEA (gene set enrichment analysis) results using the correlation levels of individual genes with TMB are shown. The number of genes in the Gene Ontology (SIZE) and other results are shown as output of GSEA. Enrichment score (ES) and normalized ES (NES) are shown with significance levels. Significance level of zero indicates P< 0.001.

**S2 Table. Molecular functions enriched for low TMB.** Preranked GSEA results using the correlation levels of individual genes with TMB. Similarly shown as with S1 Table.

S3 Table. Correlation table of DDR gene expression and mutation signature levels.

S4 Table. Correlation table of DDR gene methylation and mutation signature levels.

